# Photoreceptor calyceal processes accompany the developing outer segment, adopting a stable length despite a dynamic core

**DOI:** 10.1101/2023.10.04.560921

**Authors:** Maria Sharkova, Gonzalo Aparicio, Constantin Mouzaaber, Flavio R Zolessi, Jennifer C Hocking

## Abstract

Vertebrate photoreceptors detect light through a large cilium-based outer segment, which is filled with photopigment-laden membranous discs. Surrounding the base of the outer segment are microvilli-like calyceal processes (CPs). While CP disruption has been associated with altered outer segment morphology and photoreceptor degeneration, the role of the processes remains elusive. Here, we used zebrafish as a model to characterize CPs. We quantified CP parameters and report a strong disparity in outer segment coverage between photoreceptor subtypes. CP length is stable across light and dark conditions, while heat shock inducible expression of tagged actin revealed rapid turnover of the CP actin core. Detailed imaging of the embryonic retina uncovered substantial remodeling of the developing photoreceptor apical surface, including a transition from dynamic tangential processes to vertically-oriented CPs immediately prior to outer segment formation. Remarkably, we also found a direct connection between apical extensions of the Müller glia and retinal pigment epithelium, arranged as bundles around the ultraviolet sensitive cones. In summary, our data characterize the structure, development, and surrounding environment of photoreceptor microvilli in the zebrafish retina.

## INTRODUCTION

Microvilli extend from the apical cell surface as finger-like protrusions supported by a core of filamentous actin (F-actin) (Nambiar et al., 2010). In the small intestine and renal proximal convoluted tubule, thousands of microvilli together form a brush border, thereby massively increasing the surface area of the cell for transport of solutes between the lumen and intracellular space (Crawley et al., 2014; Coudrier et al., 1988). Sensory cells can also extend microvilli, although of varying morphologies and purposes. Best studied are the stereocilia of the inner ear hair cells, which contain thick actin bundles and are arranged in rows of increasing heights (Tilney et al., 1992; Barr-Gillespie, 2015). Stereocilia are deflected upon auditory or vestibular stimulation, leading to the opening of gated ion channels, cell depolarization, and activation of the associated sensory nerve.

Light sensation by retinal photoreceptors is mediated by the outer segment (OS), an enlarged and modified microtubule-based cilium packed with photopigment-laden membranous discs (Goldberg et al., 2016). The base of the OS is surrounded by a ring of microvilli known as calyceal processes (CPs) and presumed to have a supportive, non-sensory role. CPs extend from the apical surface of the inner segment (IS), which houses organelles such as the mitochondria and endoplasmic reticulum and performs the metabolic functions of the cell.

Although first described in the 19^th^ century, the functions of CPs remain uncertain (Schultze, 1872). CPs are found in a wide range of species, including fish and humans (Nagle et al., 1986; Sahly et al., 2012). Certain rodents such as mice and rats lack CPs altogether or possibly have a single large “tongue-like” CP or a few vestigial protrusions (Sahly et al., 2012; Volland et al., 2015). CPs house an actin core that is continuous with rootlets extending deep into the IS and, at least in some cases, anchoring at the outer limiting membrane (OLM), the location of junctions between Müller glial processes and photoreceptor ISs (Nagle et al., 1986; Williams et al., 1990).

Rod photoreceptors, responsible for vision in dim light, have a rod-shaped OS where the discs are discrete units fully enclosed within the plasma membrane (Goldberg et al., 2016). In the OSs of cones, which mediate high-acuity colour vision, the discs are lamellae continuous with one another and the plasma membrane of the IS. Photoreceptors are long-lived cells that in humans cannot regenerate. Nevertheless, the burden of oxidative damage is mitigated by the continuous turnover of the OS through creation of new discs on the basal side and removal of old discs at the apical tip through phagocytosis by the adjacent retinal pigment epithelium (RPE). One proposed function of CPs is as a barrier to restrain the growth of nascent discs (Schietroma et al., 2017). Indeed, disruption of CPs was previously associated with the overgrowth of basal discs in rods.

The significance for CPs in supporting vision was highlighted by their association with Usher syndrome, the most common form of inherited combined hearing and vision loss (Sahly et al., 2012). USH type 1 (USH1) is characterized by severe congenital hearing loss and prepubertal onset of retinitis pigmentosa (El-Amraoui and Petit, 2014). The hearing deficits caused by lack of USH1 proteins are well understood, with each contributing to the structure and function of inner ear stereocilia, but the retinal manifestations are less clear, largely because the mouse mutants do not exhibit vision problems. It was proposed that USH1 visual deficits are a result of disrupted CPs, which would explain the lack of a mouse phenotype. Indeed, it was demonstrated that CPs in frogs and macaque express all six USH1 proteins: the adhesion proteins cadherin-23 (USH1D) and protocadherin-15 (pcdh15/USH1F), the scaffolding proteins harmonin (USH1C) and sans (USH1G), the actin-bundling protein espin (USH1M), and the cytoskeletal motor protein myosin 7a (USH1B) (Sahly et al., 2012). Functional data is however limited. Primarily, morpholino knockdown of Pcdh15 in *Xenopus tropicalis* and *pchd15b* mutation in zebrafish each resulted in disrupted CPs and disorganized OSs (Schietroma et al., 2017; Miles et al., 2021).

Zebrafish have been widely adopted as a model for studying the visual system (Noel et al., 2022). The zebrafish retina exhibits the same layered organization as the human retina, except for the lack of a central fovea, and contains a mix of rods and cones (*≈*60% cones in adults (Zang and Neuhauss, 2021)). As vision-dependent predators, zebrafish use blue cones, red/green double cones, and ultraviolet-sensitive (UVS) cones for a wide spectrum of colour vision. Further, the zebrafish photoreceptors are arranged in a highly organized mosaic pattern (Raymond et al., 1995).

Here, we characterize the CPs of zebrafish photoreceptors and surrounding structures as a basis for future research into CP function. CP dimensions were analyzed across photoreceptor subtypes, with observed differences in length, width, and percent coverage of the OS. CP length is stable between light and dark conditions despite changes to height of the IS, while the actin core undergoes constant renewal. During development, photoreceptor precursors feature dynamic tangential processes that remain after differentiation. In addition, a unique actin dome structure was observed in the nascent IS, expanding above the OLM and serving as a platform for growing CPs. Finally, our data suggest a surprising interaction between apical processes of Müller glia and the RPE.

## RESULTS

### Basic CP parameters in 1 mpf zebrafish

By one month post fertilization (1 mpf), zebrafish rods and cones are functional and exhibit well-developed morphology; this time point was therefore chosen to perform a basic characterization of zebrafish CPs. First, we measured CP length in confocal images of eye cryosections stained with phalloidin conjugate to visualize F-actin (Fig. 1A,C). As the actin bundles that form the CP cores extend from roots emerging deep in the IS, we used the presence of horizontal F-actin fibers visible at the IS/OS junction, just above the mitochondrial cluster, to demarcate the IS/OS boundary. Double cones were highlighted by the zpr1 antibody and peanut agglutinin (PNA), blue cones by the anti-blue opsin antibody, UVS cones by the *sws1:GFP* transgene, and rods by the *rho:eGFP* transgene. CP length in rods was 5.9+/-0.6 μm, in double cones 3.2+/-0.3 μm, in blue cones 5.7+/-0.3 μm, and in UVS cones 6.6+/-0.9 μm. Interestingly, plotting of CP length relative to OS length revealed that CPs of rods and double cones exhibit *≈*30% OS coverage, whereas the blue and UVS cone OSs are almost completely enveloped by CPs (*≈*70–80%) (Fig. 1D). Next, CP number was counted in sagittal sections of *Tg(rho:eGFP)* retinas; 13+/-0.9 CPs were observed around blue cone OS and 20+/-0.9 CPs around double cones (Fig. 1B). Photoreceptor type was identified based on position within the photoreceptor layer, unique OS shape, and exclusion of rods (labeled by GFP). Rod and UVS cone phalloidin staining was substantially weaker and did not allow for consistent assessment. When analyzing transmission electron microscopy (TEM) sections (Fig. 1E,F,F’ and Fig. S1A,B), double cone CP diameter was significantly larger (149+/-23 nm) than of UVS cones and rods (122+/-18 nm and 131+/-17 nm, respectively), which may account for the difference in actin staining.

**Figure 1.**
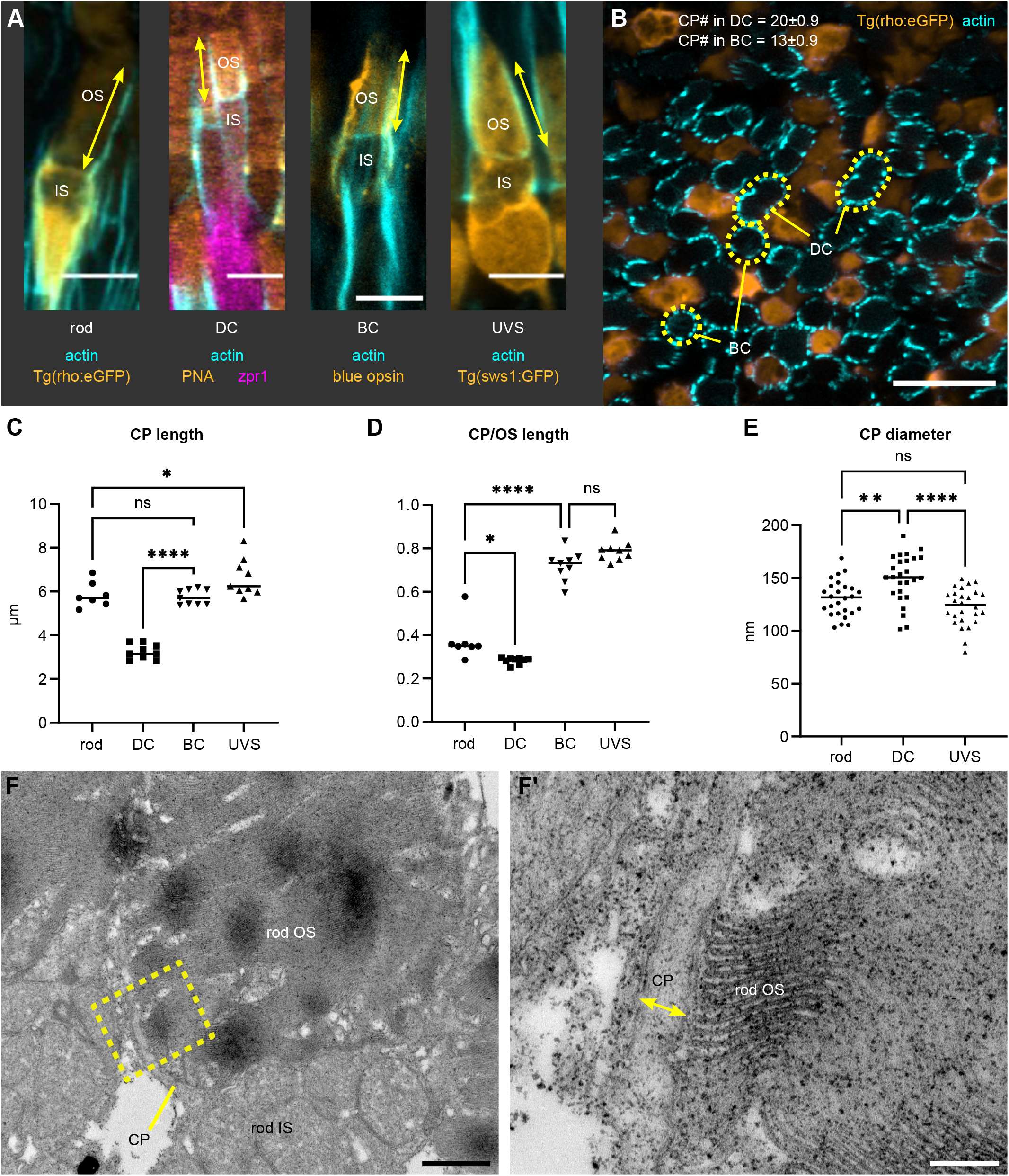
CP parameters for photoreceptor subtypes in the juvenile zebrafish retina. (A) Confocal images of 1 mpf retina stained with phalloidin (cyan). Dark-adapted rod (DA *Tg(rho:eGFP)*), light-adapted double cone (DC) (LA wild-type (WT) stained with PNA and zpr1), blue cone (BC) (LA WT stained with anti-blue opsin), and UV-sensitive cone (UVS) (LA *Tg(sws1:GFP)*); CP length is indicated by arrows. (B) Sagittal section through a 1 mpf DA *Tg(rho:eGFP)* retina (rods in orange) labeled with phalloidin (cyan). BC and DC OSs are outlined. For CP number, median and standard deviation are shown; n=5 fish. The first two graphs display CP length (C) and CP length relative to the OS length (D) for LA DC, LA BC, LA UVS cones, and DA rods; number of fish n=9 (DC, BC, UVS), n=7 (rod). (E) Graph showing CP diameter measured in rods, DC, and UVS cones in TEM images of 1 mpf WT retina, with individual measurements plotted; number of fish n=5. Statistics (C–E): median is shown; one-way ANOVA with Tukey’s test; ns—p>0.05, *— p<0.05, **— p<0.01, ****— p<0.0001. (F) Example of TEM imaging used for measuring CP diameter. Lower magnification image showing rod OS, IS, and a CP, with a yellow contour indicating the area in (F’), where CP diameter is labeled. Scale bars: 5 μm (A), 10 μm (B), 1 μm (F), 200 nm (F’).

Together, these data demonstrate that CP parameters can vary and suggest potentially different roles depending on the photoreceptor subtype.

### CP length is constant during photoadaptation

Teleosts undergo retinomotor movements as an adaptation to light conditions (Burnside and Nagle, 1983). In the dark, rod ISs shorten to bring the rod OSs closer to any incoming light, while cone ISs elongate to move their OSs further into the RPE layer. The opposite occurs in the light. Additionally, RPE melanosomes translocate into the apical RPE processes in the light and retract into the cell body in the dark.

Previously, it was demonstrated that CPs in isolated green sunfish rods shorten upon light adaptation (Pagh-Roehl et al., 1992). To compare CP length in dark-adapted (DA) and light-adapted (LA) 1 mpf zebrafish, we first assessed whether retinomotor movements can already be observed at this stage, as formerly shown for double cones (Hodel et al., 2006). We measured the distance between the OLM and IS/OS junction in rods, as well as in double, blue, and UVS cones (Fig. 2A,B) and found a significant difference between the DA and LA state for all four photoreceptor subtypes (Fig. 2C). As expected, rod ISs were longer in LA conditions, whereas double, blue, and UVS cone ISs were longer in DA zebrafish. The difference in length was most pronounced and observable in rods. Surprisingly, when we measured the length of CPs for the four photoreceptor subtypes, there was no significant difference between the DA and LA state (Fig. 2D). The LA rod CPs were mostly obscured by the pigment granules in RPE villi: therefore *crystal* zebrafish lacking pigment in the eye were used to measure rod CP length (Antinucci and Hindges, 2016). Since the lack of pigment could influence photoreceptor health, we compared DA rod CP length in *Tg(rho:eGFP)* and *crystal* zebrafish. There was no significant difference between the two groups (Fig. S2A,B), demonstrating that *crystal* fish are an appropriate model for analysis.

**Figure 2.**
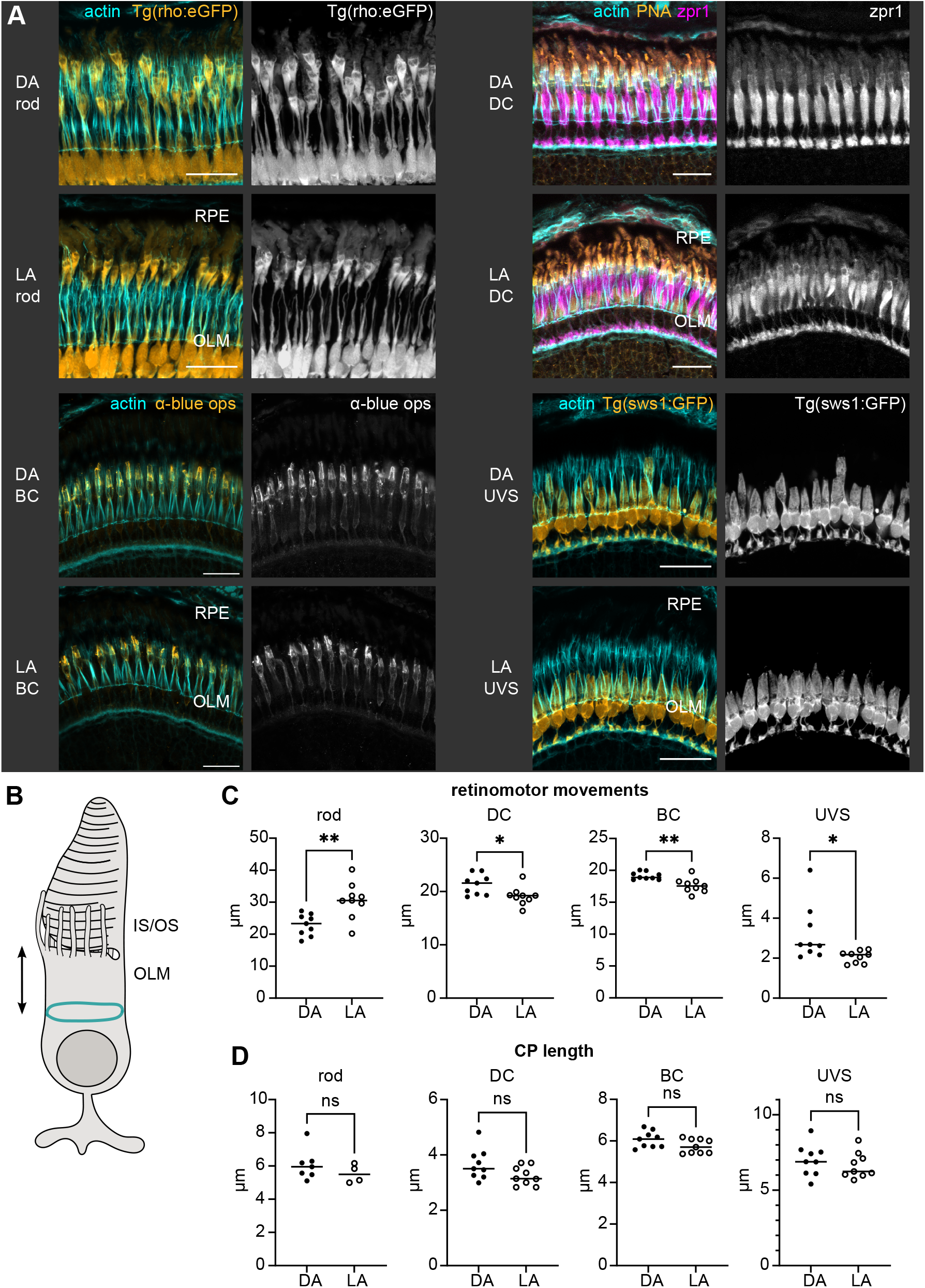
Retinomotor movements and CP length in dark-adapted and light-adapted 1 mpf zebrafish retina. (A) Confocal images of 1 mpf DA and LA outer retina sections stained with phalloidin (cyan). From left to right, top to bottom: rods (*Tg(rho:eGFP)*), double cones (DC) (WT stained with PNA and zpr1), blue cones (BC) (WT stained with anti-blue opsin), UV-sensitive cones (UVS) (*Tg(sws1:GFP)*). Scale bars: 20 μm. (B) Schematic depiction of measurement for the IS–OLM distance. (C) The graphs show the extent of cellular retinomotor movements as a distance between the apical IS and the OLM in each photoreceptor cell type, DA versus LA state. (D) Graphs displaying the CP length in photoreceptors in DA vs LA fish. Statistics: number of fish n=9 (rods, DC, BC, UVS), n=7 (rod DA CP length), n=4 (rod LA CP length); median is shown; unpaired t-tests with Welch’s correction; two-tailed p-value; ns—p>0.05, * — p<0.05, **— p<0.01.

The data we obtained indicate CP length remains constant while ISs undergo retinomotor movements, implying CPs could have a stabilizing role to support OS translocation.

### CP precursors emerge prior to OS development

Previous scanning electron micrographs of the chicken and *Xenopus* retina suggest that CPs emerge from the apical IS before OS appearance (Olson, 1979; Sahly et al., 2012; Wai et al., 2006). In addition, they appear to undergo a selection, where some microvilli are eliminated as a cilium emerges from the IS (Olson, 1979). To investigate early OS and CP development in zebrafish photoreceptors, TEM imaging was performed. When inspecting 70 hpf (hours post fertilization) eyes, several stages characterized by location and distinct morphology were observable within each retina. Peripheral photoreceptors were at an early stage of differentiation with no evidence of the apical mitochondrial clustering characteristic of the IS. The apical cell surfaces of the photoreceptors and RPE here were flat, creating a smooth interface between the two cells (Fig. 3A). Some of the peripheral photoreceptors exhibited processes on their apical surfaces, but these extended tangentially, parallel to the photoreceptor layer, rather than extending towards the RPE (Fig. 3A, arrowheads). Interestingly, RPE cells also appeared immature, with only a few pigment granules positioned between photoreceptors and the RPE nuclei. When moving away from the periphery and towards the central retina, the interface between the photoreceptors and RPE now appeared rougher, with multiple apical protrusions visible on the surface of each cell type and interdigitating with each other (Fig. 3B, arrowheads). At the same time, the photoreceptor apical domain expanded to form the IS, becoming filled with clustering mitochondria. Further, RPE granules increased in number. Still, most photoreceptors lacked a budding cilium. In areas closer to the ventronasal patch and dorsocentral region, where cells differentiate earliest (Schmitt and Dowling, 1999), we observed large ISs with dense mitochondrial clusters and newly forming OSs bordered by CPs (Fig. 3C, arrow). In addition, some cells had a cilium emerging from the IS surface. The nascent photoreceptor cilia were swollen, as previously documented (Nilsson, 1964b), and grew directly into the RPE layer, such that the cilium was enveloped by an RPE pocket. Interestingly, the cilia appear to penetrate the RPE layer alone as no CPs were detected within the RPE pocket prior to the formation of OS discs (Fig. 3C and Fig. S1C).

**Figure 3.**
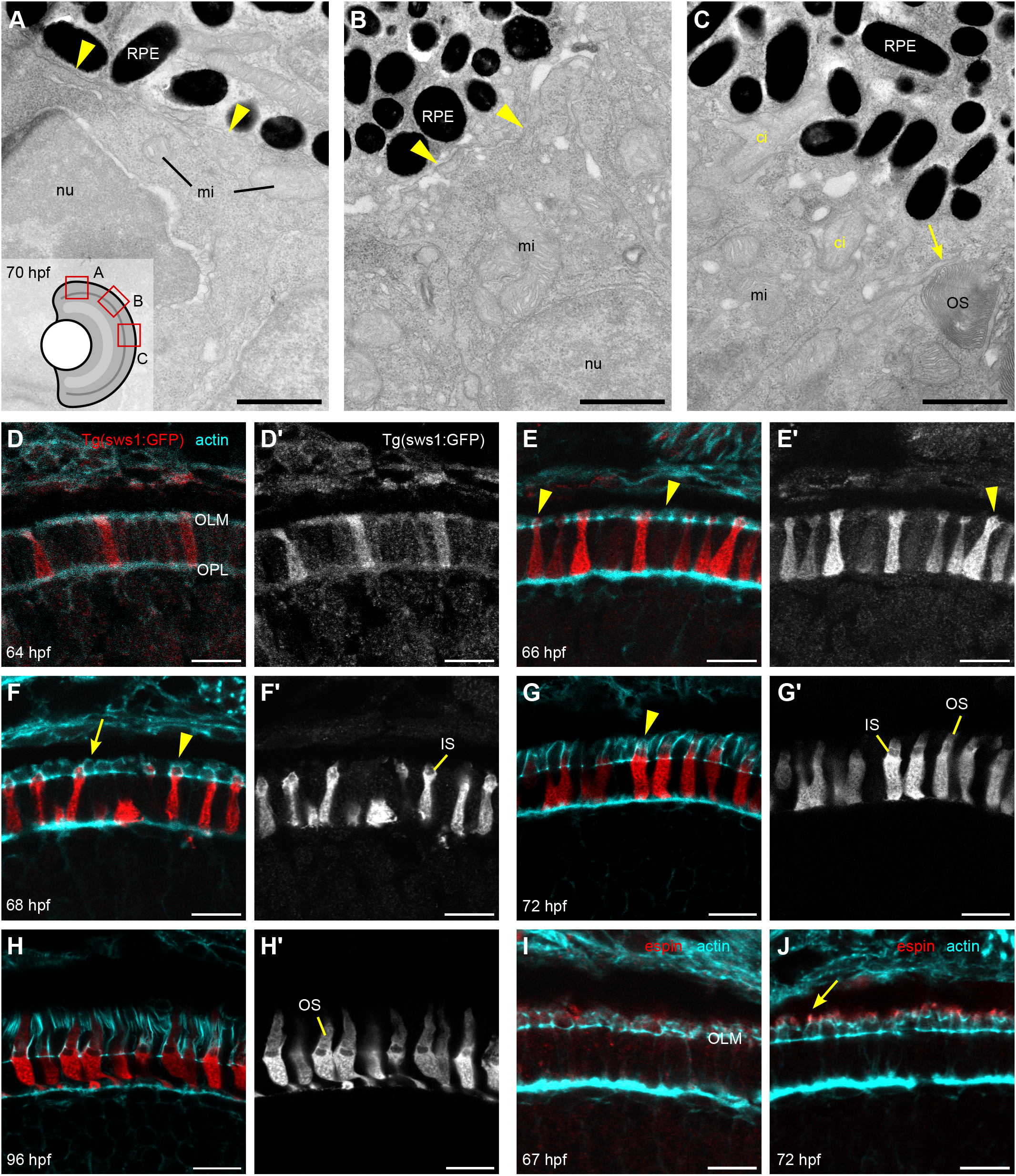
Details of IS, OS, and CP development in zebrafish retina. (A–C) TEM micrographs depicting the progression of IS and OS development in 70 hpf WT embryonic retina. A small schematic inset in A shows the approximate position of each panel. (A) Peripheral retina, the photoreceptor/RPE interface is flat, with an isolated apical photoreceptor process extending parallel to the interface, as indicated by arrowheads. mi - mitochondria; nu - nucleus. (B) When moving away from the periphery, processes (arrowheads) can be observed emerging from both the RPE and photoreceptor apical surfaces, creating an interdigitating IS/RPE interface. (C) Dorsocentral region, one photoreceptor has an OS with well-developed discs and a visible adjacent CP (arrow), while two other photoreceptors are at the emerging cilium (ci) stage. (D–H) Confocal images of *Tg(sws1:GFP)* (red) outer retina sections at 64, 66, 68, 72, and 96 hpf stained with phalloidin (cyan). (D) Early photoreceptors with a columnar morphology and an actin-rich apical domain, but no distinct IS. (E) Arrowheads pointing at actin dome-like structure in the IS; (E’) filopodia emerging from IS apical surface (arrowhead). (F) Different IS/actin dome shapes: round and rectangular (arrow and arrowhead, respectively). (G) Arrowhead indicates CPs. (I,J) Confocal images of 67 and 72 hpf outer retina sections stained with phalloidin (cyan) and anti-espin (red). (J) An arrow highlights espin localization to the CPs in 72 hpf fish. Number of fish analyzed n=3 (A–C), n=7 (D), n=5 (E), n=8 (F,G), n=6 (H), n=4 (I,J). Scale bars: 1 μm (A–C), 10 μm (D–J). 12

To obtain further detail about F-actin distribution during photoreceptor development, we performed confocal imaging of zebrafish ocular cryosections stained with phalloidin. *Tg(sws1:GFP)* embryos were selected for sectioning because UVS cones are the earliest forming photoreceptors within the zebrafish retina and the transgene provides clear visualization of the cells (Crespo and Knust, 2018). As expected, different morphological stages were observable within a single section due to the wave-like development of photoreceptors over time across the retina (Raymond et al., 1995). For consistency, we analyzed only the dorsocentral retina in Fig. 3D–J. At 64 hpf, very few ISs were observed and most photoreceptors, including UVS cones, featured a flat, actin-rich apical domain (Fig. 3D). A broad expansion of the IS occurred around 66 hpf, with mitochondria beginning to cluster apically, as indicated by the region of weak GFP signal (Fig. 3E). The nascent IS also featured F-actin extending above the OLM in a dome-like shape (Fig. 3E, arrowheads) and filopodia-like projections emerging from the apical surface of some photoreceptors (Fig. 3E’, arrowhead). At 68 hpf, further IS elongation occurs and a mitochondrial cluster is clearly delineated (Fig. 3F’, arrow). Some photoreceptors at this stage retain the rounded apical surface of the IS (Fig. 3F, arrow), while others have now assumed a rectangular shape (Fig. 3F, arrowhead). Faintly stained vertical projections sprouting from the IS actin dome are observed in many cells. We expect that most cells have developed a small OS or at least a cilium by 68 hpf; however, this was difficult to observe, likely owing to interference by the pigment of the adjacent RPE pocket. In the 72 hpf retina, further IS elongation has occurred and short actin-filled processes surrounded a well-formed UVS OS, now visible (Fig. 3G, arrowhead). By 96 hpf, photoreceptors exhibit further OS and CP growth, as well as changes to synaptic morphology (Fig. 3H).

CPs undergo an initial growth phase between 68 and 72 hpf. To better understand the transition from precursors to CPs in zebrafish embryos, we analyzed the localization of espin (USH1M), an actin bundling protein associated with microvillar growth in other cell types (Desban et al., 2019). At 67 hpf, espin is weakly expressed within the IS actin dome, above the OLM (Fig. 3I). Remarkably, espin strongly localizes to the nascent processes, suggesting an active bundling phase coinciding with CP growth (Fig. 3J and Fig. S2C).

CPs accompany the OS from an early stage, yet are not associated with the nascent cilium. Remarkably, photoreceptor microvilli exist prior to the cilium or OS appearing, and the IS actin dome precedes OS formation and serves as a base for CP sprouting.

### Tangential processes persist during photoreceptor differentiation

When analyzing photoreceptor development prior to OS formation, we captured apical processes of diverse morphology — tangential processes on the progenitors and vertical processes atop the nascent ISs. To identify whether the two represent different stages of a single structure or develop individually, we applied mosaic labeling obtained by injecting a DNA construct driving the expression of a membrane form of GFP under the *crx* promoter region (*crx:EGFP-CAAX*), highlighting the external shape of isolated cells. At the periphery of the cell differentiation area in 72 hpf embryo retinas, photoreceptors and progenitors at different stages can be found. For example, some cells exhibit the typical shape of early photoreceptor progenitors, with a rounded cell body (i.e., has not yet elongated along the apical-basal axis) and a profusion of thin cellular processes extending radially from the edges of the apical cell surface (Fig. 4A, Movie S1). These processes have been previously described at earlier developmental stages as “tangential processes” (Aparicio et al., 2021) and they are characterized by highly dynamic behavior, evident upon time-lapse observation of GFP-positive cells at the periphery of 60–72 hpf retinas (Fig. 4B, Movie S2).

**Figure 4.**
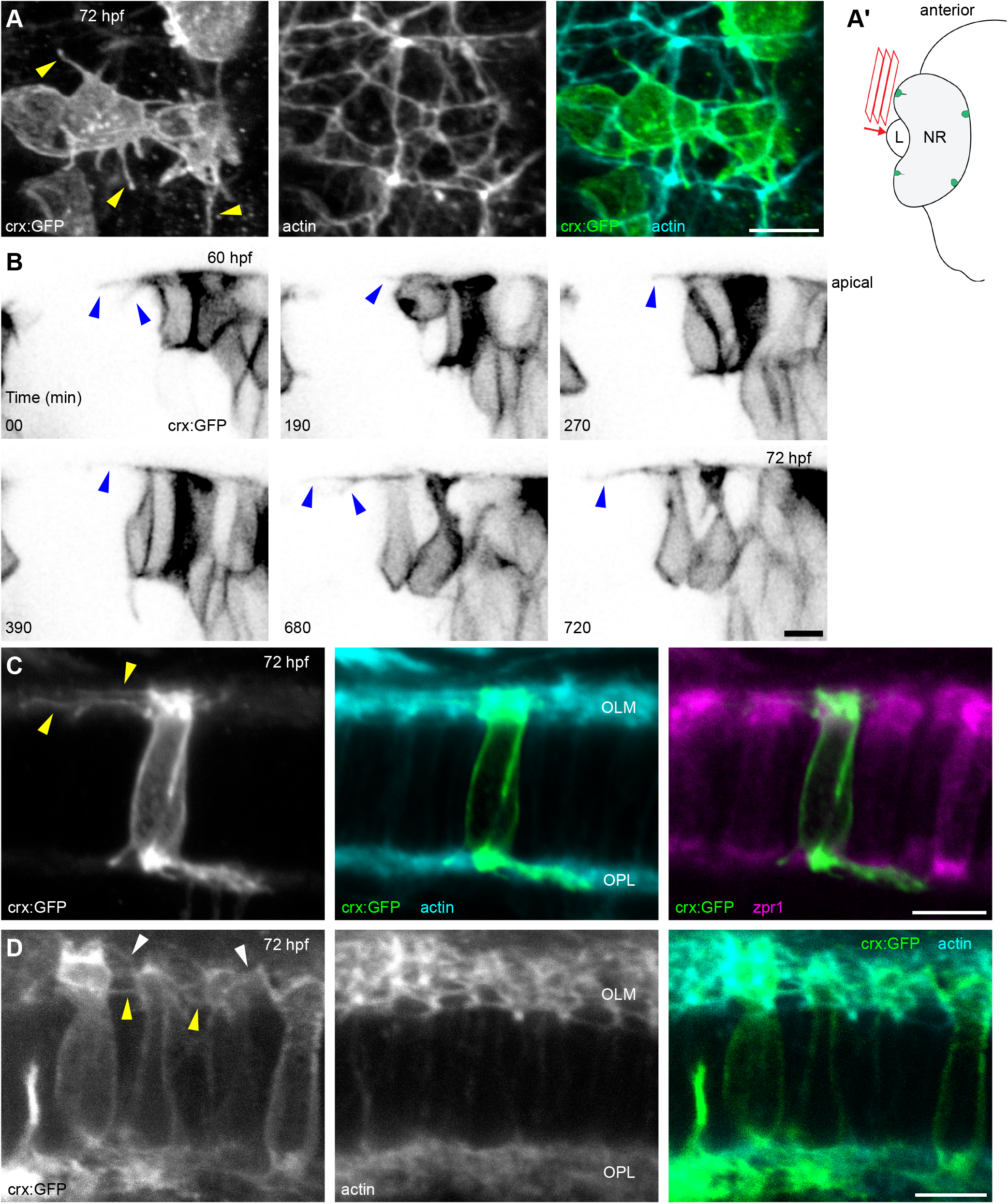
Tangential processes on photoreceptor progenitors and differentiating photoreceptors in the 72 hpf retina. (A) Apical view of the peripheral-most *crx:EGFP-CAAX* (*crx:GFP*) expression area, showing a few photoreceptor progenitors profusely extending tangential processes (arrowheads). F-actin staining with TRITC-phalloidin highlights the sub-apical adhesion rings. (A’) Diagram depicting the orientation of acquisition in (A); L — lens, NR — neural retina. (B) Time-lapse experiment of *crx* :GFP-injected embryos, showing a peripheral area of the retina displaying photoreceptor progenitors extending highly dynamic tangential processes on the apical surface (arrowheads). (C) Early differentiating photoreceptor, displaying long tangential processes (arrowheads). (D) Differentiating photoreceptors at the IS-forming stage, showing short processes extending from this expanding apical membrane. Some of the processes are connected at the OLM, indicating they might be tangential processes (yellow arrowheads), while others originate at more apical positions and extend in different directions (white arrowheads). Number of embryos analyzed n=8 (A,C,D), n=4 (B). Scale bars: 5 μm.

Around the same region, other cells with *crx* promoter-driven GFP expression have already acquired an apico-basally elongated conformation, indicating they are post-mitotic differentiating photoreceptors (Fig. 4C,D). Interestingly, we observed zpr1 antibody-labeled cells, just at the onset of IS formation and still harboring relatively long tangential processes (Fig. 4C). Other photoreceptors, more advanced in the differentiation process and showing an evident forming IS, display many cell processes extending from their apical portion, albeit shorter than at earlier stages (Fig. 4D). Some of these processes originate from the interphase between the cell body and the IS, at the level of the OLM, at the same position and direction as earlier tangential processes. Some others, however, originate from the apical dome above the OLM and extend in various directions. The tangential processes are lost in subsequent stages, as evident from images of the *Tg(sws1:GFP)* transgenic line in Fig. 3. Cells with ISs and tangential processes are visible in the dorsocentral retina at 66 hpf (Fig. 3E’), but the tangential processes are absent by the time the OS has formed at 72 hpf (Fig. 3G’). Movie S3 and Fig. S3 document the transition away from long, dynamic tangential processes as photoreceptors mature and begin to form specialized apical regions.

In summary, we discovered a brief period of overlap between tangential processes and the onset of CP formation, coinciding with the emergence of the IS. While tangential processes briefly coexist with CPs on developing photoreceptors, the two types of actin-based cellular protrusions are dynamically and morphologically distinct.

### CPs feature a dynamic actin core

Intestinal brush border microvilli exhibit rapid actin recycling through growth of the filaments at the microvillar tips and disassembly inside the cell body, a process known as treadmilling (Meenderink et al., 2019). On the other hand, hair cell stereocilia in the ear feature only tip turnover, with the shaft remaining stable for months (Zhang et al., 2012; Narayanan et al., 2015; Drummond et al., 2015). To determine which type of actin dynamics is characteristic for CPs, we used Tol2 transgenesis to create fish carrying a random insertion containing the heat shock promoter *hsp70l*, zebrafish *actb1* cDNA, and a myc tag (Fig. 5A). The construct included a *cmlc2:EGFP* transgenesis marker to drive GFP expression in the heart and allow for selection of positive embryos. 24 hours after heat shock, the fish were euthanized and processed for microscopy. In 6 dpf (days post fertilization) injected larvae featuring mosaic myc expression, newly introduced tagged actin was observed at the OPL (weak expression) and in CPs and their roots, with particularly strong expression in the latter (Fig. S2E). The OLM was almost entirely devoid of tagged actin, although strongly stained by phalloidin. No positive cells were detected in the control zebrafish. For further analysis, a stable transgenic line was generated (referred to as *Tg(hsp:act-myc)*). Only a few photoreceptors with low baseline actin-myc expression were occasionally observed in control zebrafish at 1 mpf (Fig. 5B). In contrast, all zebrafish in the heat shock group had strong actin-myc expression in the majority of cone photoreceptors (Fig. 5C). The localization of tagged actin in cones of the juvenile fish was similar to that observed in injected larvae: absent at the OLM, diffuse in the synaptic layer and the IS, and highly concentrated in CPs and CP roots (Fig. 5D,D’). Occasionally, a rod IS not entirely concealed by the RPE was detected, always myc-positive (Fig. 5D’, arrow).

**Figure 5.**
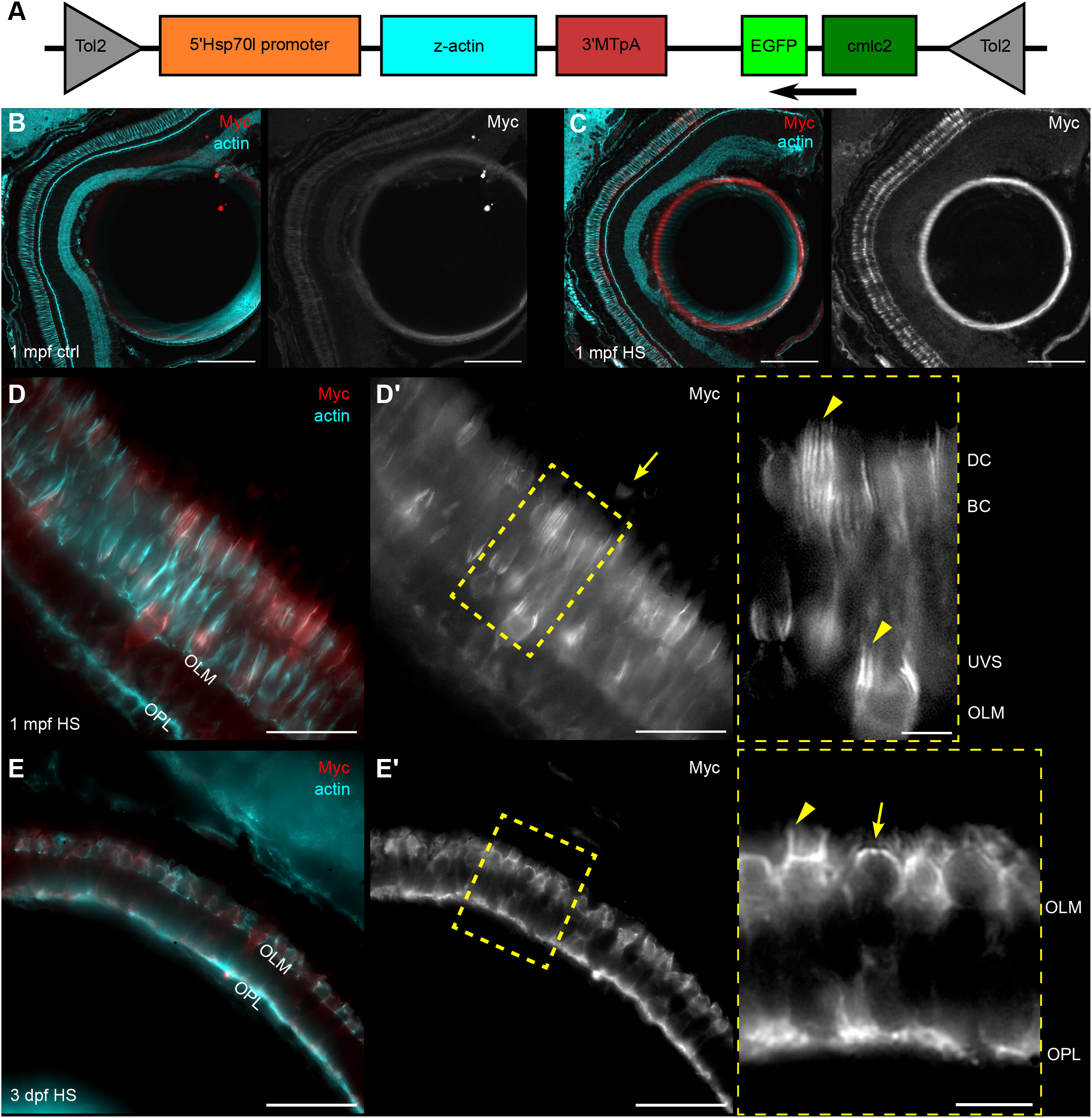
Induced actin is incorporated into CP cores in zebrafish. (A) Graph representing various components of the construct injected into 1-cell stage WT embryos. (B–E) Micrographs of *Tg(hsp:act-myc)* zebrafish retina stained with phalloidin and anti-myc antibody. (B) Control 1 mpf *Tg(hsp:act-myc)* eye. (C) Eye of 1 mpf *Tg(hsp:act-myc)* fish 24 h after heat shock (HS). (D) Higher magnification of a photoreceptor layer of heat shock treated 1 mpf fish; arrow in (D’) points at the rod IS; inset shows enlarged yellow box contents from (D’), arrowheads highlight myc localization to CPs. (E) 3 dpf *Tg(hsp:act-myc)* embryo 6 h after heat shock. (E’) Yellow box indicates position of enlarged area in the inset; actin-myc expression in the IS actin dome (arrow), and in the CPs (arrowhead). Number of fish analyzed n=11 (B), n=12 (C), n=7 (D), n=11 (E). Scale bars: 100 μm (B,C), 20 μm (D–E), 5 μm (insets).

To determine actin dynamics while CPs are extending alongside the growing OSs, 3 dpf *Tg(hsp:act-myc)* embryos were euthanized 6 hours after heat shock. Again, only a few positive cells were detected in control eyes (Fig. S2F). In the heat shock group, the photoreceptor layer exhibited strong actin-myc expression (Fig. 5E,E’). Compared to 1 mpf photoreceptors, there was stronger myc labeling at the OPL, and an occasional weak signal at the OLM was observed. Both CPs and the roots featured high actin-myc incorporation, in contrast to espin localizing mostly to CPs at this stage (Fig. 3J). Also of note, actin-myc could be observed throughout the IS actin dome of immature photoreceptors in the peripheral retina.

Despite maintaining a consistent length during retinomotor movements, actin cores of photoreceptor microvilli and their IS roots undergo constant incorporation of new actin monomers in both embryonic and juvenile fish.

### Complexity of structures organizing the OS layer

Photoreceptor OSs are encased in a supportive environment that includes CPs, a complex interphotoreceptor matrix, and extensive RPE villous protrusions (Ishikawa et al., 2015; Steinberg et al., 1977). Less recognized are processes extended by Müller glial cells. Above the OLM, Müller glia extend microvilli and, at least in zebrafish, also longer, thicker apical processes that reach UVS cone OSs (Zou et al., 2012). Given that glial and RPE processes protrude into the relatively constricted space between the bulky photoreceptor OSs, the possibility arises that they not only interact with the photoreceptors, but also with each other. To visualize the positioning and complexity of these support arrangements, we labeled retinal sections from 1 mpf *Tg(gfap:GFP)* zebrafish, in which Müller glia express GFP and the full cell morphology can be well visualized. The long apical glial protrusions colocalized with phalloidin staining of thick actin bundles and extended alongside UVS cone OSs all the way to the tips (Fig. 6A, arrowhead and Fig. S2D). Incredibly, the apical glial processes overlapped, in very close proximity, with the RPE villi visualized by zpr2 antibody and descending towards the OLM (Fig. 6A,C) (Hanovice et al., 2019). In a tangential view, the phalloidin-stained thick actin bundles within the long glial processes are visible surrounding the UVS cone OSs, at a ratio of five glial processes per OS. Further, the actin bundles are adjacent to rod ISs, together forming a regular pattern as part of the zebrafish photoreceptor mosaic (Fig. 6B,D). Notably, Müller glial apical processes do not protrude beyond the OLM in 3 dpf embryonic retina (Fig. S2G), and therefore do not accompany the emerging OS.

**Figure 6.**
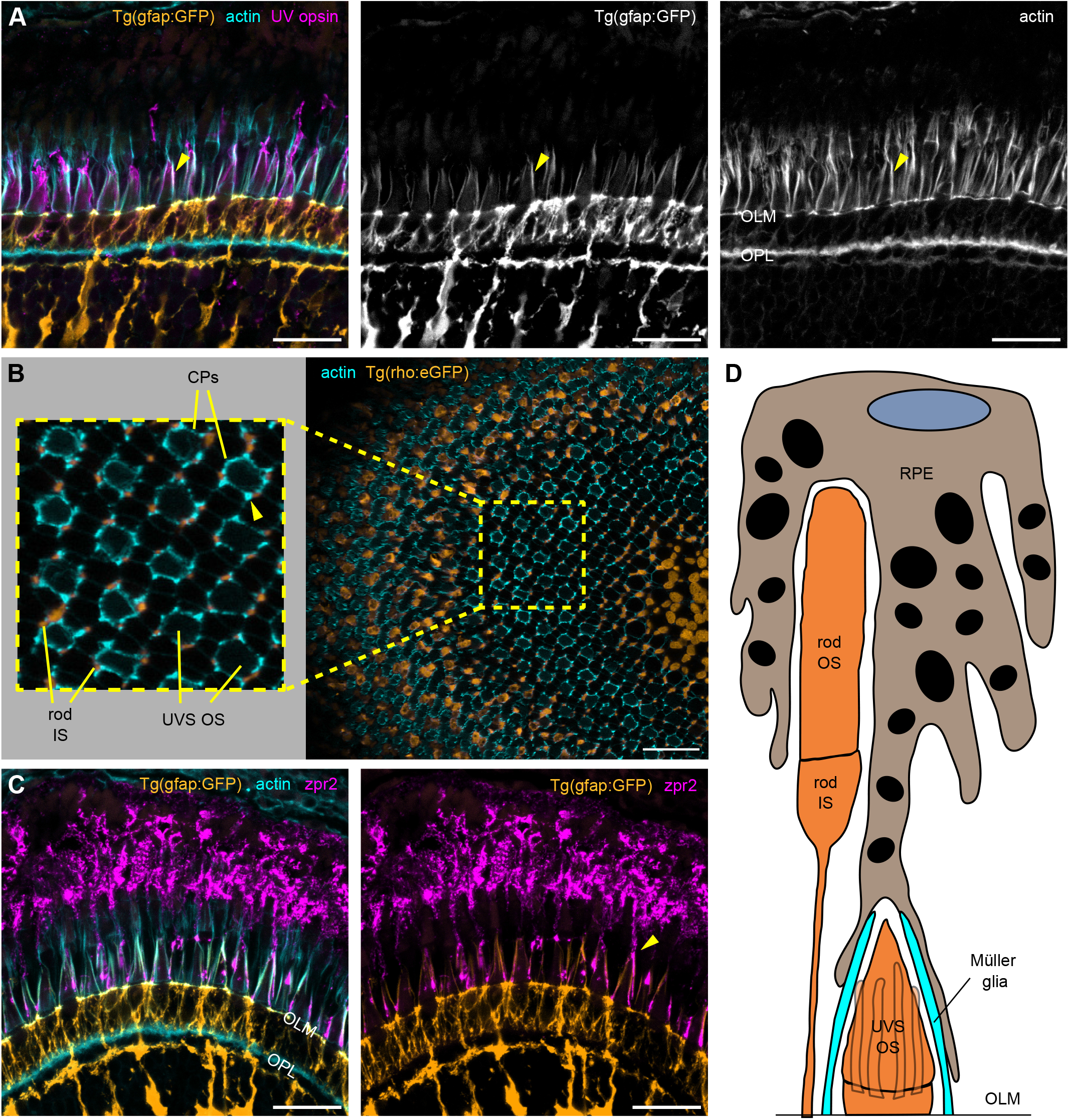
Zebrafish Müller glia and RPE protrusions enclose UVS cones OSs. (A–C) Confocal images of 1 mpf zebrafish retina sections incubated with phalloidin (cyan) and UV opsin or zpr2 antibody (magenta). (A) *Tg(gfap:GFP)* zebrafish with Müller glia cell bodies highlighted by GFP (orange) show long glial processes above the OLM stretching alongside UVS cone OSs and colocalizing with thick actin bundles (arrowheads). (B) Sagittal section through *Tg(rho:eGFP)* retina with an enlarged area demonstrating rod ISs (orange) adjacent to thick actin bundles (arrowhead) surrounding UVS cones OSs. (C) RPE apical villi, stained with zpr2 antibody, extend towards the OLM and localize in close proximity to the apical Müller glia processes, as observed in *Tg(gfap:GFP)* retina. (D) A model illustrating the organization of supporting cells in the photoreceptor layer. UVS cones feature both Müller glia and RPE protrusions around the OS. Number of fish analyzed n=3 (A,C), n=5 (B). Scale bars: 20 μm (A–C).

Müller glia and RPE represent the two main cell types supporting the homeostasis of photoreceptors. We demonstrate that their apical protrusions overlap to create a unique encapsulation of UVS cone OSs.

## DISCUSSION

While CPs remain poorly understood, a possible association with the retinal USH1 phenotype brought them to attention as a potentially critical aspect of photoreceptor biology (Sahly et al., 2012; Schietroma et al., 2017; Miles et al., 2021). As zebrafish is a favourable model for photoreceptor disease studies (Noel et al., 2022), our detailed examination of CP characteristics in wildtype zebrafish will provide a useful reference for future investigation. Most notably, we characterized the transition from dynamic tangential processes to vertical CPs just prior to OS formation, as well as how CPs undergo continuous turnover of their actin cores while maintaining constant lengths.

### Assessment of zebrafish CP parameters

We characterized basic parameters of CPs in zebrafish using confocal microscopy and TEM. While images of zebrafish CPs were previously shown in a TEM analysis of photoreceptors (Tarboush et al., 2012) and in the context of the *pcdh15b* mutation (Miles et al., 2021), our data provide a quantitative and detailed assessment of CPs in relation to the various photoreceptor subtypes. Comparing our findings to green sunfish, another teleost species where data is available, zebrafish CPs are in the same length range (3–6 μm vs 5 μm in green sunfish) but fewer in number (12–14 in zebrafish blue cones vs 23–26 in green sunfish single cones) (Nagle et al., 1986). When considering mammalian species with quantification data available, the number of CPs extended by zebrafish blue cones is comparable to that of macaque cones (14–16) (Sahly et al., 2012). In addition, macaque cone CP length is similar to our measurements in zebrafish double cones (3 μm), though diameter is larger (244 nm vs 150 nm in zebrafish). Notably, zebrafish CPs are longer than intestinal microvilli (1– 3 μm), but similar to renal microvilli (3–5 μm) and the microvilli of cerebrospinal fluid-contacting neurons (3–4 μm) (Sharkova et al., 2022). As reported for other species (Sahly et al., 2012; Schietroma et al., 2017), we observed substantial differences in basic parameters between photoreceptor subtypes, not only between rods and cones, but also between short and long cones.

The most surprising finding in analysis of CP length was the *≈*70–80% coverage of blue and UVS cone OSs given that CPs are always described as encircling the base of the OS. Of note, blue and UVS cone OSs are closest to the OLM and most distant from the RPE. It is plausible that the extended CPs compensate for diminished support from apical RPE processes and help guide the translocating OS lamellae.

Still, RPE villi do extend alongside the UVS OS and feature extensive overlap with Müller glial apical protrusions. This implies a special regulation of UVS OS dynamics and a potential direct interaction between the RPE and Müller glia. While the two cell types are both well characterized as supportive of photoreceptor function, they are typically portrayed as physically separate in the literature. Indeed, we found only one reference, from 1964, of contact between RPE and Müller glial processes in the bullfrog, *Rana Pipiens* (Nilsson, 1964a). Better acknowledged is evidence of RPE-derived factors being necessary for the proper functioning of the Müller glia (Jablonski et al., 2001). RPE signaling was also demonstrated to induce Müller glia proliferation both in vitro (Jaynes and Turner, 1995; Goczalik et al., 2005) and in vivo (Webster et al., 2019). Contacts between glia and RPE processes could play an important role in maintaining photoreceptor health and function, and may have been overlooked in other species.

Retinomotor movements are a feature of teleost and amphibian retinas and we examined whether the contraction/elongation of the ISs was associated with changes in CP length. Surprisingly, we detected no difference in CP length between light and dark conditions. Having confirmed that retinomotor movements occur by this point (1 mpf) (Hodel et al., 2006), we therefore expect CPs to maintain constant length in older fish as well. Notably, our results differ from previous experiments on green sunfish showing light-induced contraction of rod CPs occurring alongside elongation of ISs (Pagh-Roehl et al., 1992). This may be a species difference, although only rods were examined in the sunfish. Interestingly, retinomotor movements are not an entirely actin-driven process, as microtubule translocation plays a role at least in the elongation of cone myoids, suggesting a mechanism for decoupling CPs from IS movements (Lewis et al., 2018; Burnside, 1976). Microtubules are abundant in both rod and cone ISs (Verschueren et al., 2022).

Of note, there are contrasting views regarding UVS cone participation in retinomotor movements (Menger et al., 2005; Neuhauss, 2010). Our data support the idea that UVS cone ISs change length upon light adaptation, albeit to a lesser extent than those of rods and double cones. Interestingly, retinomotor movements are also not equal across all cells. For example, light-adapted shortened rods are divided into two rows (also previously described in (Pagh-Roehl et al., 1992)), and the dark-adapted UVS cone row features isolated individual cells that are noticeably longer than the majority.

### CP development: before and after the OS

Neuroepithelial progenitors undergo considerable morphological change during their development into photoreceptors. Our goal here was to learn more about how CPs fit into the context of photoreceptor maturation.

Several papers described the presence of processes atop the IS prior to OS emergence. In scanning electron microscopy images of *Xenopus* photoreceptors, the developing CPs appeared on the apical surface of the IS (Sahly et al., 2012). Initially immature, they change their morphology after OS emergence. Similarly, two papers examining chick retina showed abundant microvilli emerging from the “ball-like” ISs as they bulged above the OLM (Olson, 1979; Wai et al., 2006). The microvilli protruded both vertically and laterally, without any overt organization. Previous work also showed the presence of very dynamic filopodia-like tangential processes emerging from the edges of the apical surface of differentiating zebrafish photoreceptors, though well before IS expansion (Aparicio et al., 2021). Here, we observed the presence of both vertical (CP precursors) and lateral (tangential) processes prior to OS formation. While the IS expands, tangential processes undergo retraction and CPs emerge, and we observed a brief period of processes extending in multiple directions, suggesting a dramatic change in actin dynamics at the apical cell surface. Further, a primary cilium is present on the apical surface during the transition from neuroepithelial cell to photoreceptor, but appears to be retracted before newly emerging as the nascent OS (Aparicio et al., 2021). Importantly, we observed that CPs, while present, do not abut the newly formed cilium. Instead, the cilium is fully encased within the RPE, and contact with CPs only begins once the first discs are formed.

The apical dome formation just prior to OS and CP emergence was demonstrated previously for chick and zebrafish photoreceptors (Olson, 1979; Wai et al., 2006; Crespo and Knust, 2018). A similar structure has not been described for maturing renal epithelial cells, cerebrospinal fluid-contacting neurons, or inner ear hair cells just prior to microvilli formation and therefore the actin dome is likely related more to IS maturation or OS formation than to CP emergence (Desban et al., 2019; Barr-Gillespie, 2015; Gaeta et al., 2021). Indeed, the clustering of mitochondria in the apical portion of the cell is an early indicator of the specialization of the apical photoreceptor region and occurs concomitantly with the formation of the actin-lined dome. The IS subsequently transitions from a dome shape to a cylindrical shape as the OS begins to form discs and becomes encircled by CPs.

Differentiating cerebrospinal fluid-contacting neurons adopt a circumferential apical actin ring from which grow the actin bundles giving rise to the microvillar cores (Desban et al., 2019). Photoreceptors have a similar actin ring at the OLM, which is maintained from the junctions between neighboring neuroepithelial cells (Spitznas, 1970). The actin lining the apical dome is anchored at the OLM, as are the roots for the nascent CPs. Notably, F-actin remains at the IS/OS junction of mature photoreceptors, visible as a line immediately above the mitochondrial cluster.

Our data align with previous findings showing that photoreceptor microvilli change form over the course of development, as illustrated by Fig. 7; however, we discovered a surprising and distinct transition from tangential, dynamic filopodia to vertical, static microvilli.

**Figure 7.**
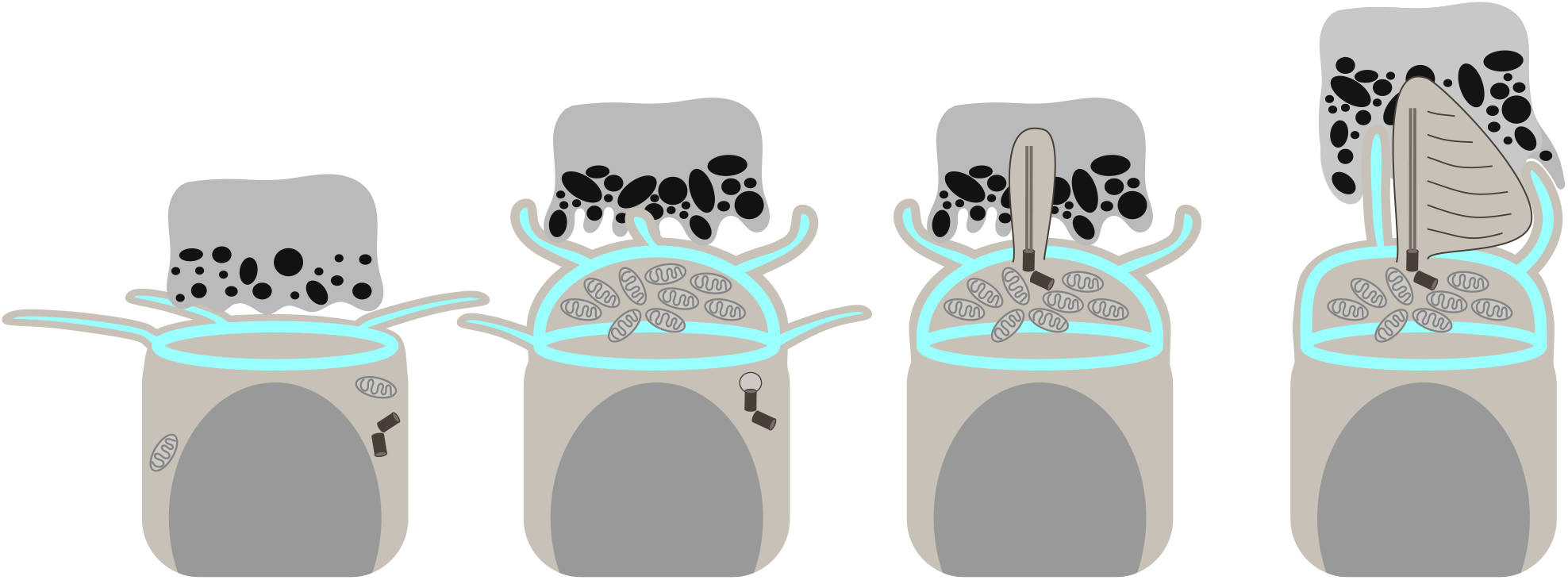
Diagram depicting stages of photoreceptor CP, IS, and OS development in embryonic zebrafish. From left to right: CPs, IS, and OS of zebrafish photoreceptors undergo distinct alterations during development. First on the left: no distinct IS is observed; photoreceptors feature tangential processes apically, an actin ring at the OLM, and flat RPE/IS interface. Next, the IS becomes prominent, outlined by an actin dome, and vertical processes (presumably CP precursors) appear, while the RPE/IS interface becomes rougher. Tangential processes originating near the OLM area are retained. Further, a cilium, the future OS, emerges and enters the RPE pocket, with no processes adjacent to it. Finally, the cilium starts generating discs, the CPs associate with the new OS, and the IS becomes more rectangular in shape. Please note that the diagram does not accurately depict relative sizes of photoreceptors and RPE in order to highlight the apical region of the former.

### Implications of dynamic CPs

In this paper, we provide insight into actin dynamics of photoreceptors. CPs and their IS roots feature fast incorporation of new actin monomers, whereas actin associated with the cell-cell junctions at the OLM and in the OPL synapses are relatively stable in juvenile zebrafish. There was no visible difference in localization of induced actin between cone subtypes. In the developing 3 dpf retina, both the CP cores and the IS actin dome appear highly dynamic, as anticipated based on the rapid morphological changes we observed in those structures. The synapses also demonstrate a high expression of induced actin, possibly coinciding with their maturation (Schmitt and Dowling, 1999). On the other hand, the OLM is stable at 3 dpf, showing limited incorporation of new actin.

The speed of microvillar actin turnover varies depending on the type of cell, with two models being particularly well-researched: rapid treadmilling in brush border microvilli (Loomis et al., 2003) and tip turnover on a stable shaft in stereocilia (Zhang et al., 2012; Narayanan et al., 2015; Drummond et al., 2015). Stereocilia are neuronal microvilli and share a set of basic actin cross-linkers with photoreceptor CPs (espin, fascin, and fimbrin/plastin), and the processes were reported to express the Usher complex proteins known to create links between the stereocilia (Sahly et al., 2012; Lin-Jones and Burnside, 2007; Höfer and Drenckhahn, 1993; McGrath et al., 2017; Schietroma et al., 2017; Verschueren et al., 2022). However, stereocilia have a mechanosensory role supported by thick actin bundles and a unique staircase arrangement (Tilney et al., 1980). Indeed, despite the Usher-inspired comparison of CPs to stereocilia, the former exhibit actin dynamics resembling the brush border.

Treadmilling involves the addition of actin monomers to the F-actin plus ends at the microvilliar tips and removal from the cytosolic minus ends. Using our heat shock system, we showed rapid turnover in CPs but could not elucidate the exact pattern of actin monomer addition and removal. However, the actin bundle in CPs is reportedly oriented as in other microvilli, with the plus ends at the distal tip, suggesting a similar mechanism of actin renewal (O’Connor and Burnside, 1981; Pagh-Roehl et al., 1992).

Our data shows that CPs maintain a constant length despite continual renewal of their actin cores. The consistency of CP length and OS coverage within each photoreceptor subtype but disparity between subtypes suggests careful regulation of CP growth. However, we do not yet understand the function of CPs or the importance of precise length control. One proposed CP function is to restrain growth of nascent discs at the basal OS. Indeed, overgrowth of photoreceptor discs was observed when proposed CP–OS linker proteins, Pchd15 or Cdh23, were reduced or absent (Miles et al., 2021; Schietroma et al., 2017). Alternative CP functions could be to provide general structural support for the OS, possibly in conjunction with surrounding tissues, or transport metabolites to the OS, bypassing the connecting cilium. Despite being discovered more than 150 years ago and residing adjacent to the cellular compartment where vision begins, CPs remain a mystery. Further research is necessary to uncover their role in photoreceptor biology.

## MATERIALS AND METHODS

### Zebrafish husbandry

Zebrafish were handled at the University of Alberta aquatic facilities according to standard protocols and with ethics protocol approved by Animal Care and Use Committee (AUP1476) and at the Zebrafish Laboratory, Institut Pasteur de Montevideo, following the approved local regulations (CEUA-IPMon, and CNEA). Embryos were collected from a breeding and raised in embryo medium (1x E2, Zebrafish International Resource Center (ZIRC)) at 28.5°C with a 14/10 hours light/dark cycle. At 5–6 dpf, larvae were transferred to the aquatic facility. Zebrafish were euthanized using an overdose of methanesulfonate salt (Acros Organics, pH adjusted to 7.0).

TL and AB zebrafish were used as wild-types, with only one line used throughout an experiment. *Crystal* zebrafish lacking pigment in the eye and the body was generated in the laboratory of Dr. Ted Allison based on the previously described *crystal* line (Antinucci and Hindges, 2016; Balay, 2018). Transgenic strains were used to examine UV cones (*Tg(sws1:GFP)*, (Takechi et al., 2003)), rods (*Tg(rho:eGFP)*, (Hamaoka et al., 2002)), Müller glia (*Tg(gfap:GFP)*, (Bernardos and Raymond, 2006)), and tagged actin incorporation after heat shock (*Tg(hsp:act-myc)*, see details below).

### Light adaptation setup

For experiments with LA vs DA comparison, DA zebrafish were kept in the dark overnight + 1 hour. The other group was LA for 1 hour, and both groups were euthanized at the same time in the morning. The DA fish were handled under dim red light.

### Tissue preparation and immunostaining

Whole euthanized zebrafish were fixed with 4% paraformaldehyde (PFA) in phosphate-buffered saline (PBS) overnight at 4°C. The fixative was washed out with PBS in three washing steps. Afterwards, a 17.5% sucrose solution was added until the fish sank (from *≈*1 hour for 3 dpf embryos to 1 day for 1 mpf juveniles). They were then left in 35% sucrose at 4°C overnight. Next, the fish were oriented in plastic cryomolds filled with optimal cutting temperature compound (Tissue-Tek, Sakura Finetek). The blocks were frozen on dry ice and stored at -80°C until sectioning. 12 μm sections were cut with the Thermofisher Shandon E, Leica CM1520, or Leica CM1900 cryostat. The sections were transferred onto Superfrost Plus slides (Fisherbrand) and kept at -20°C.

After warming up the slide for 5 minutes, the tissue area was outlined with a lipid pen, followed by a short rinse with PBS. Next, the sections were permeabilized with PDT (0.1% Triton X-100, 1% dimethyl sulfoxide in PBS). The sections were blocked for 1 hour with 5% goat or donkey serum in PDT (depending on the secondary antibody type) and subsequently incubated with primary antibodies diluted in blocking solution at 4°C overnight. Next, secondary antibodies and conjugated phalloidin diluted in blocking solution were added for 1 hour at room temperature. All antibodies and conjugated dyes are listed in Table S1. After washing, the slides were mounted with mowiol-based homemade mounting medium (pH=8.5, RI*≈*1.51, 2.5% DABCO), coated with coverslips, and kept at 4°C.

For visualization of tangential processes, embryos were grown in 0.003% phenylthiourea (PTU, Sigma), fixed overnight at 4°C by immersion in 4% paraformaldehyde in phosphate buffer saline, pH 7.4 (PFA-PBS). For whole-mount immunostaining, all subsequent washes were performed in PBS containing 1% Triton X-100.

### Generating *Tg(hsp:act-myc)* line

To generate *Tg(hsp:act-myc)* zebrafish, we followed the Tol2kit protocol combining Gateway recombination technology and Tol2 transposon-based incorporation (Kwan et al., 2007). To obtain zebrafish actin (zact) cDNA (transcript actb1-201, ENS-DART00000054987.7), mRNA was isolated from 3 dpf TL embryos (RNeasy, Qiagen; RNAlater, Invitrogen), and AffinityScript QPCR cDNA Synthesis Kit (Agilent Technologies) with a set of specific primers (F: CCATGGATGAGGAAATCGCTG; R: AGAAGCACTTCCTGTGGACGATG) was applied. All primers were ordered from Integrated DNA Technologies as 25 nmole oligos with standard desalting purification. For higher yield, we cloned the zact sequence into the bacterial plasmid pCR 2.1 (TOPO TA cloning kit with One Shot TOP10 chemically competent cells, Invitrogen). TOP10 chemically competent cells were also used in other steps.

Next, zact sequence was amplified with primers containing attB sites (F: GGGGACAAGTTTGTACAAAAAAGCAGGCTC-CATGGATGAGGAAATCGCTG; R: GGGGACCACTTTGTACA-AGAAAGCTGGGTAGAAGCACTTCCTGTGGACGATG) using a high-fidelity polymerase (Phusion, NEB). To create a middle entry clone pME-zact, we performed a BP reaction cloning attB-zact product into a donor vector pDONR221 (BP Clonase II, Invitrogen, 11789020). The subsequent LR reaction (LR Clonase II, Invitrogen, 11791) combined three entry clones and one destination vector (p5E-*hsp70l* + pME-zact + p3E-MTpA + pDestTol2CG2) into one construct (pDestTol2CG2; *hsp70l*:zact-MTpA).

On the morning of injection, Tol2 mRNA and the construct (final concentration 25 ng/μL each) were combined and 1 nL of the mixture was injected into 1-cell stage TL embryos. Positive embryos were selected at 1 dpf based on the presence of GFP signal in the heart. Injected fish were grown into adulthood and incrossed; positive embryos from this breeding were used in heat shock experiments. Additionally, a group of injected fish underwent preliminary heat shock experiments to confirm that myc-tagged actin is properly expressed after heat shock and to test various heat shock conditions.

### Generating *crx* mosaic embryos

pDestTol2pA2;crx:EGFP-CAAX (Aparicio et al., 2021), together with Tol2 transposase mRNA were injected into the one-cell stage *roy* background (Ren et al., 2002) according to standard techniques.

### Heat shock

1 mpf juvenile zebrafish were transferred into a separate tank with a Hygger Titanium Aquarium Heater (HG-802, 50W), heat shocked at 39°C and euthanized 24 hours later. Embryos and larvae were subjected to 37°C in a water bath and euthanized 6 hours after that. In both cases, the duration of the heat shock was 1 hour. Control zebrafish were transfered into a tank/tube with same parameters, but without heating.

### Fluorescent imaging and processing

For the confocal imaging, Zeiss LSM510, LSM700 or LSM800 microscopes were used with a 63x 1.4 NA oil objective. Zeiss Elyra 7 Lattice SIM was used to visualize fine detail of tagged actin incorporation presented in Figure 5(D,E). ZEN (2009, 5.5 & 3.0 black for Elyra), ImageJ (1.54f), and Imaris (9.8.2) were used to process the images.

### TEM

Zebrafish were fixed with a mixture containing 2.5% glutaraldehyde and 2% PFA diluted in 0.1 M phosphate buffer. After three wash steps, the post-fixation was achieved with 1% osmium tetroxide to provide contrast for the sample. The fish were washed again, gradually dehydrated with ethanol, and infiltration with Spurr’s resin was performed overnight. Next, they were embedded in flat molds containing fresh resin and left in the oven at 70°C overnight. The blocks were cut at the ultramicrotome into 70–90 nm sections that were stained with uranyl acetate and lead citrate.

The images were acquired with the Philips/FEI (Morgagni) Transmission Electron Microscope with Gatan Camera operating at 80 kV. TEM images were processed in ImageJ (version 1.54f).

### Time-lapse imaging

Embryos were selected at 50–60 hpf, anesthetized using 0.04 mg/mL MS222 (Sigma), and mounted in 1% low melting-point agarose, containing 0.003% N-phenylthiourea and 0.04 mg/ml MS222/tricaine (Sigma) over n° 0 glass bottom dishes (MaTek). During overnight image acquisitions, embryos were kept in Ringer’s solution (116 mM NaCl, 2.9 mM KCl, 1.8 mM CaCl2, 5 mM HEPES pH 7.2) with 0.04 mg/mL MS222. Live acquisitions were made using a Zeiss LSM 880 laser confocal microscope with a 40x 1.2 NA objective and glycerol:water (75:25) immersion medium. Stacks around 40 μm thick were acquired in bidirectional mode, at 1 μm spacing and 512 × 512 pixel resolution every 10 min.

### Image analysis

The sample size was calculated using the Boston University resources (URL: www.bu.edu/research/ethics-compliance/animal-subjects/animal-care/research/sample-size-calculations-iacuc/, last accessed on 2024-02-26). To perform all statistical tests and to create graphs, GraphPad Prism software (9.5.0) was used. CP number and the TEM data were assessed in one eye of the fish. For all other experiments, both eyes were analyzed and the average was calculated to represent the fish. When comparing LA vs DA zebrafish, the images were blinded.

## Footnotes

## Supporting information

Figure S1-3; captions for Movie S1-3

## Acknowledgements

We acknowledge the employees of Health Sciences Laboratory Animal Services and Science Animal Support Services, University of Alberta, and of the Zebrafish Lab, Institut Pasteur de Montevideo, for their excellent fish care. We thank the Cell Imaging Core, U. Alberta, and Dr. Kiera Smith for Elyra access and training, the Advanced Microscopy Facility, Dr. Kacie Norton for her help with the TEM sample preparation and imaging, and the Advanced Bioimaging Unit at the Institut Pasteur Montevideo for their support and assistance with the confocal microscopes. We thank Dr. Andrew Simmonds for providing mounting medium and access to Imaris, Dr. Sarah Hughes for access to the LSM700 microscope, Dr. James Bartles for sharing the espin antibody, Dr. Anastassia Voronova for access to the Thermofisher Shandon E cryostat, Dr. Qiumin Tan for access to the Leica CM1520 cryostat, Dr. Tom Hobman for the 9E10 anti-myc antibody, Dr. Ted Allison and Dr. Paul Chrystal for sharing the Tol2kit plasmids and *crystal* zebrafish line, and Dr. Edan Foley for the Tol2 transposase mRNA. Many thanks to Dr. Paul Chrystal for critically reading the manuscript.

## Data availability

All relevant data can be found within the article and its supplementary information

## Author contributions

Conceptualization: M.S., F.R.Z., J.C.H.; Methodology: M.S., G.A., F.R.Z., J.C.H; Formal analysis: M.S., G.A., F.R.Z., J.C.H; Investigation: M.S., G.A., C.M.; Resources: J.C.H., F.R.Z.; Data curation: Writing - original draft: M.S., F.R.Z., J.C.H.; Writing - review and editing: M.S., F.R.Z., G.A., J.C.H; Visualization: M.S., G.A.; Supervision: F.R.Z., J.C.H.; Project administration: F.R.Z., J.C.H.; Funding acquisition: F.R.Z., J.C.H.

