## Supplementary material for "Photoreceptor calyceal processes accompany the developing outer segment, adopting a stable length despite a dynamic core": Figure S1-3; captions for Movie S1-3

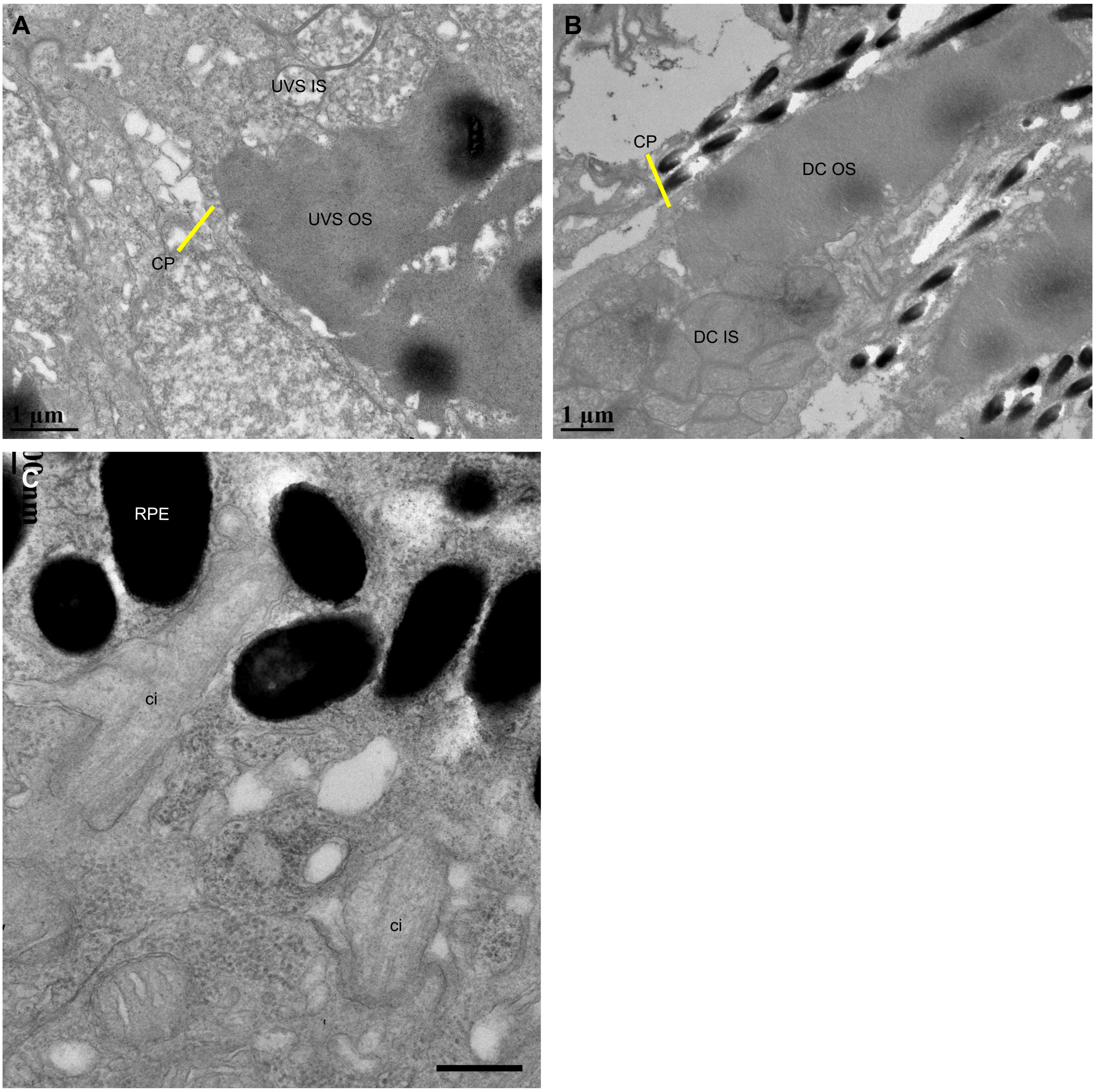


**Figure S1.** (A,B) TEM micrographs of UVS cone (A) and DC (B) with CP, IS, and OS highlighted. (C) TEM micrograph of a 70 hpf central retina, enlarged area from Fig. 3C for a closer view at nascent cilia (ci). Number of fish analyzed n=5 (A,B), n=3 (C). Scale bars: 1 µm (A,B), 0.4 µm (C).

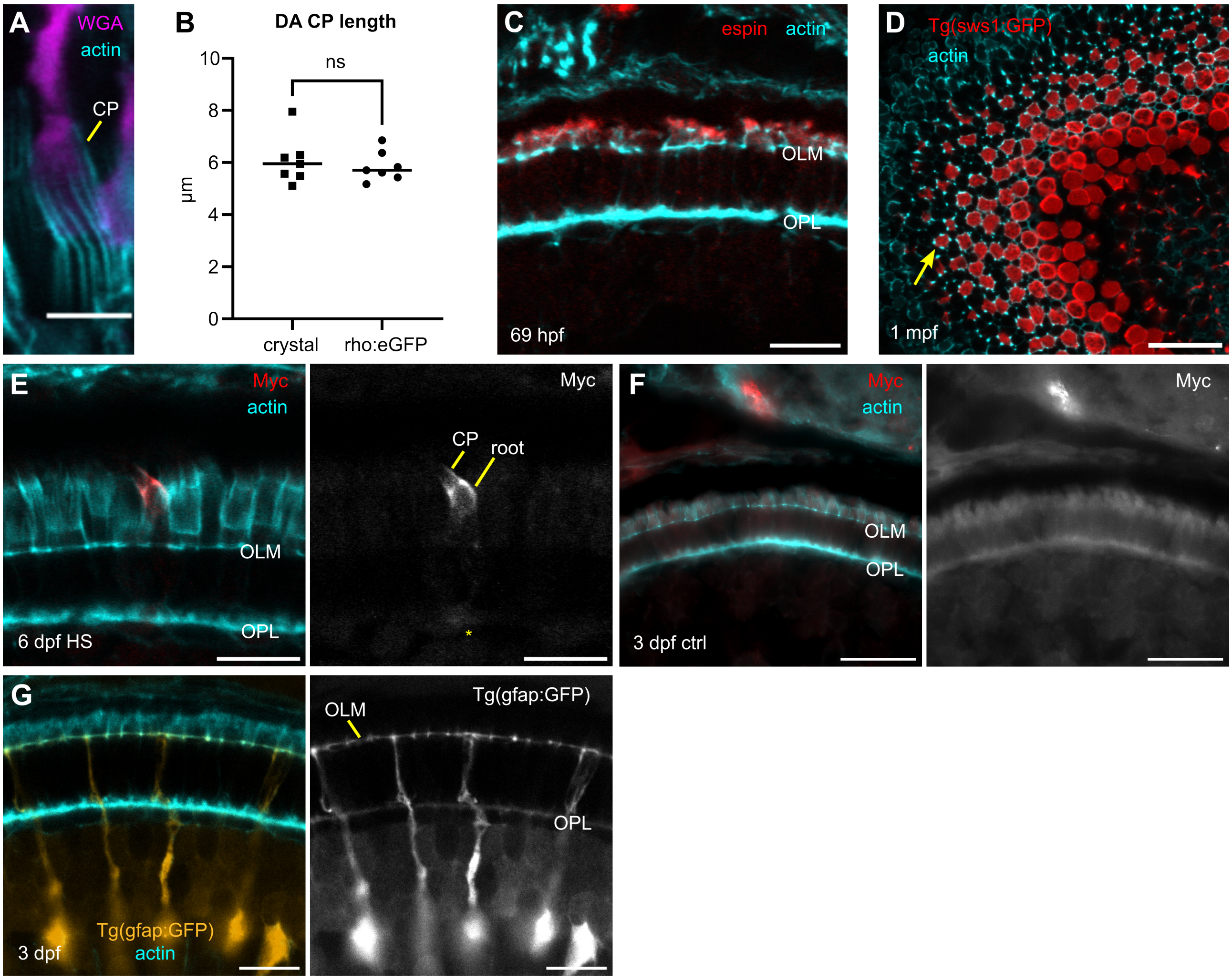


**Figure S2.** (A) Confocal image of a 1 mpf crystal zebrafish photoreceptor layer stained with WGA and phalloidin, a single rod is shown. (B) Graph displaying the comparison between CP length in 1 mpf DA *crystal* fish and *Tg(rho:eGFP)*; n=7 in both groups median is shown; unpaired t-tests with Welch's correction; two-tailed p-value; ns—p>0.05. (C) Confocal image of 69 hpf outer retina sections stained with phalloidin (cyan) and anti-espin (red). (D) *Tg(sws1:GFP)* (GFP in red) 1 mpf zebrafish retina stained with phalloidin (cyan), sagittal section. Arrow is pointing at UVS OS surrounded by thick actin bundles. (E) Confocal images of 6 dpf Tol2/hsp70l:zact-MTpA-injected zebrafish 24 h after heat shock stained with phalloidin and anti-myc antibody. Actin-myc localization in the calyceal process and its root is highlighted. Asterisk indicates diffuse staining at the OPL. (F) Control 3 dpf *Tg(hsp:act-myc)* photoreceptor layer. (G) Confocal image of *Tg(gfap:GFP)* 3 dpf zebrafish retina (GFP shown in orange) stained with phalloidin (cyan); no glial processes above the OLM are observed. Number of fish analyzed n=7 (A), n=4 (C), n=6 (D), n=8 (E), n=5 (F), n=4 (G). Scale bars: 5 µm (A), 10 µm (C,E,G), 20 µm (D,F).

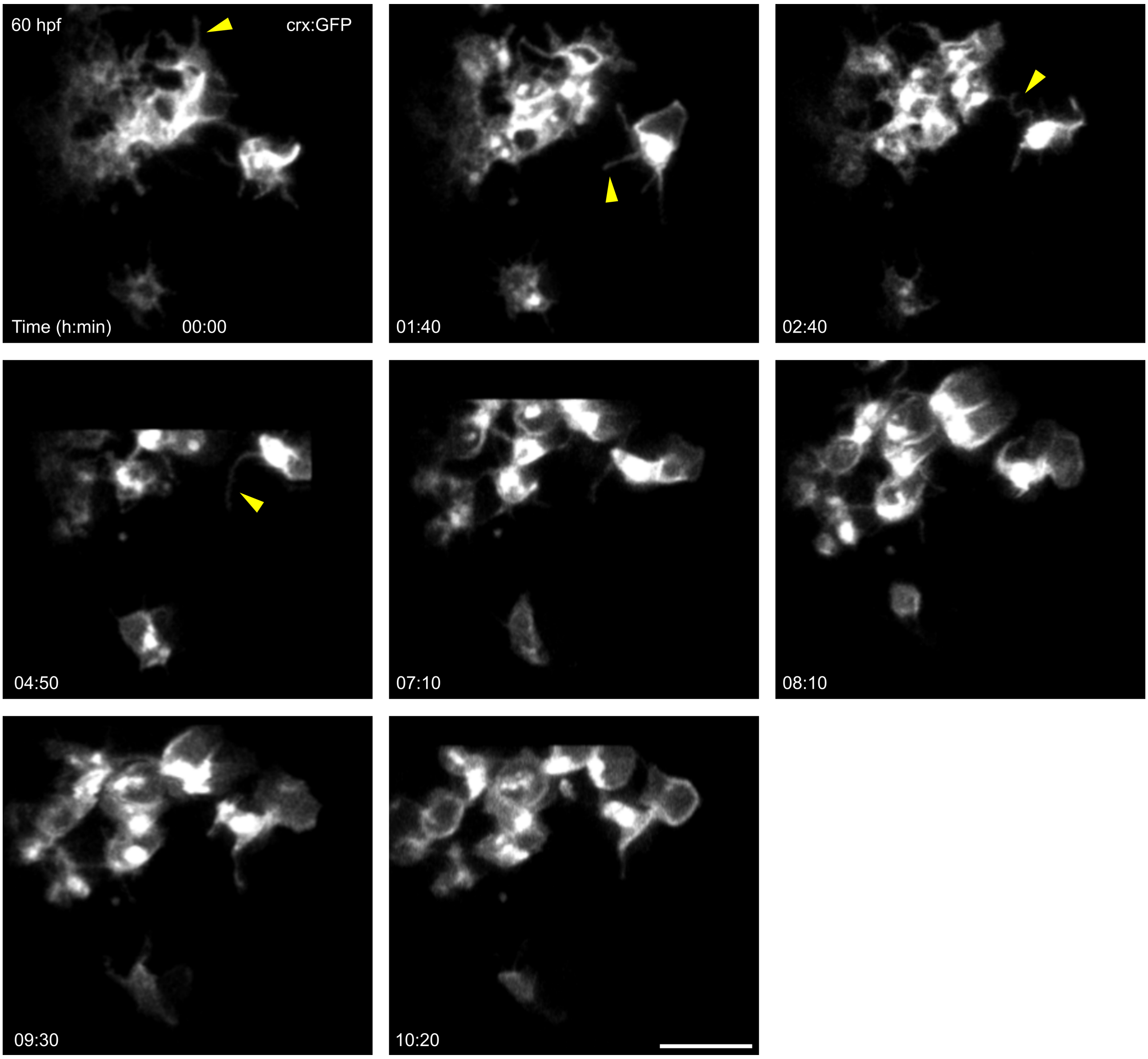


**Figure S3.** Selected time points from Movie S3. Confocal time-lapse experiment (maximum intensity projection) of a *crx:GFP* transgenic embryo from 60 hpf to 72 hpf, apical view. Arrowheads highlight dynamic tangential processes sprouting (and retracting) from the apical surface of the photoreceptor progenitors. See Fig. 4A' for orientation. Scale bar: 10 μm.

**Movie S1.** 3D reconstruction from a confocal stack at high magnification of the apical retina from a 72 hpf *crx:EGFP-CAAX* (*crx:GFP*, green) transgenic embryo, labeled with phalloidin (F-actin, cyan). Scale bar: 5 μm. See Fig. 4A for further reference.

**Movie S2.** Confocal time-lapse experiment (maximum intensity projection) of a *crx:GFP* transgenic embryo from 60 hpf to 72 hpf, with a temporal resolution of 10 min. During this time, the imaged cells at the apical-most region of the retina display the extension and retraction of tangential processes. Scale bar: 5 μm. See Fig. 4A for further reference.

**Movie S3.** Confocal time-lapse experiment (maximum intensity projection) of a *crx:GFP* transgenic embryo from 60 hpf to 72 hpf, apical view. During this time, the imaged cells display the extension and retraction of tangential processes. Temporal resolution: 10 min. Scale bar: 10 μm. See Fig. S3 for selected time points.

Table S1: List of labeling reagents for fluorescence microscopy.

| Molecule type | Host | Fluorophore | Supplier | Catalogue # |
|---|---|---|---|---|
| anti-espin antibody | rabbit | - | Bartles lab | |
| anti-GFP antibody | rabbit | - | Invitrogen | A11122 |
| anti-blue opsin antibody | rabbit | - | Kerafast | Q9W6A8 |
| anti-UV opsin antibody | rabbit | - | Kerafast | EJH013 |
| anti-myc 9E10 antibody | mouse | - | ATCC | CRL-1729 |
| zpr1 antibody | mouse | - | ZIRC | zpr-1 |
| zpr2 antibody | mouse | - | ZIRC | zpr-2 |
| anti-mouse antibody | goat | Alexa Fluor 680 | Invitrogen | A21057 |
| anti-rabbit antibody | goat | Alexa Fluor 680 | Invitrogen | A27042 |
| anti-mouse antibody | donkey | Alexa Fluor 555 | Invitrogen | A31570 |
| anti-mouse antibody | donkey | Alexa Fluor 647 | Invitrogen | A31571 |
| anti-rabbit antibody | donkey | Alexa Fluor 546 | Invitrogen | A10040 |
| anti-rabbit antibody | goat | Alexa Fluor 488 | Invitrogen | A11034 |
| phalloidin | - | CF633 | Biotium | 00046 |
| phalloidin | - | iFluor 488 | Cayman Chemical | 20549 |
| phalloidin | - | rhodamine | Abcam | ab235138 |
| phalloidin | - | TRITC | Sigma-Aldrich | P1951 |
| PNA lectin | - | Alexa Fluor 568 | Invitrogen | L32458 |
| WGA lectin | - | FITC | Sigma | L4895 |
